# AptaGPT: Advancing aptamer design with a generative pre-trained language model

**DOI:** 10.1101/2024.05.23.594910

**Authors:** Shijian Ding, Xin Yang, Chi Ho Chan, Yuan Ma, Sifan Yu, Luyao Wang, Aiping Lyu, Baoting Zhang, Yuanyuan Yu, Ge Zhang

## Abstract

Aptamers, synthetic oligonucleotide ligands, have shown significant promise for therapeutic and diagnostic applications owing to their high specificity and affinity for target molecules. However, the conventional Systematic Evolution of Ligands by Exponential Enrichment (SELEX) for aptamer selection is time-consuming and often yields limited candidates. To address these limitations, we introduce AptaGPT, a novel computational strategy that leverages a Generative Pre-trained Transformer (GPT) model to design and optimize aptamers. By training on SELEX data from early rounds, AptaGPT generated a diverse array of aptamer sequences, which were then computationally screened for binding using molecular docking. The results of this study demonstrated that AptaGPT is an effective tool for generating potential high-affinity aptamer sequences, significantly accelerating the discovery process and expanding the potential for aptamer research. This study showcases the application of generative language models in bioengineering and provides a new avenue for rapid aptamer development.

## Introduction

In recent years, aptamers, known for their unique ability to bind molecular targets selectively, have emerged as a class of molecules with far-reaching implications in medicine. These oligonucleotide ligands are often called “chemical antibodies,” they can be specifically designed to target a wide variety of molecules, including non-immunogenic molecules that are not recognized by antibodies ^1,2^. Aptamers possess structural flexibility, facile chemical synthesis, and low immunogenicity, making them optimal candidates for therapeutic and diagnostic applications in various diseases ^3^. The “Systematic Evolution of Ligand Enrichment” (SELEX) technology is the primary methodology used to identify and select aptamers^4–6^.

The SELEX process involves the construction of a library of random nucleotide sequences, their subsequent mixing with target molecules, and the isolation of the binding sequences. These sequences were then amplified, and the process was iteratively repeated to improve affinity and specificity. Following several rounds of screening, the high-affinity aptamers were cloned and serialized for further applied research^5^. Furthermore, incorporating high-throughput sequencing (HTS) technology dramatically enhances the effectiveness and precision of aptamer screening during SELEX^7,8^. HTS-SELEX allows for the quick identification of many sequences, improves sequence variability, and optimizes the screening procedure. Despite the implementation of high-throughput sequencing to enhance the efficiency of aptamer selection, it is essential to note that there are some biases in the sequencing. Moreover, the SELEX approach may have a low success rate, limiting the number of candidate aptamers that can be synthesized for subsequent affinity characterization^9^.

Numerous computational methods have been devised to analyze HT-SELEX data and identify optimal aptamers. Computational tools such as APTANI^10^, AptaCluster^11^, AptaTRACE^12^, and RaptRanker^13^ have been utilized to improve the effectiveness of aptamer screening. Nevertheless, the capacity to identify prospective aptamers is constrained by the outcomes of the SELEX experiment. To address this issue, Iwano et al. developed RaptGen^14^, a computational tool that employs a variational autoencoder (VAE)^15^ and latent representation learning, which has led to significant advances in aptamer design. It is important to note that VAE can be trained on specific datasets, such as intermediate rounds of SELEX, which may result in a limited generation quality. In contrast, generative language models can be pre-trained on a large and diverse corpus of text^16^, acquiring a wide range of language patterns and knowledge. Furthermore, this pre-training can be fine-tuned for specific tasks, enhancing the model’s overall performance^17^. This enables generative language models to fully exploit the multi-round sequence information presented in SELEX.

This study proposes an advanced computational strategy using the Generative Pre-trained Transformer (GPT)^18^ model for aptamer design. The GPT is a transformer-based deep learning model that interprets and generates complex sequence data. It has performed well in DNA^19^ and protein^20,21^ design, demonstrating its ability to generate biomolecular sequences. In this study, we selected the third and sixth rounds of SELEX data to pre-train and fine-tune the GPT model, respectively. Subsequently, computational screening methods were employed to identify candidates with a high binding capacity from the generated sequence space. These aptamers were then compared with those enriched in the 20th round to assess similarity and binding ability, further demonstrating the effectiveness of AptaGPT. In conclusion, this study demonstrates the potential of generative language models in aptamer design and provides a new pathway for improving the efficiency of SELEX.

## Results

### Overview of AptaGPT

In this study, we aimed to overcome the limitations of the traditional SELEX method by utilizing a GPT language model to design aptamers. This model utilizes unsupervised learning to process phrases or incomplete sentences from a vast corpus as input. It can effectively forecast subsequent text and produce coherent text based on the initial input structure. This approach facilitates the intelligent analysis of the evolutionary traits of aptamer sequences and proficiently generates a new set of sequences. Here, we focused on B-cell maturation antigen (BCMA) and performed 3, 6, and 20 rounds of enrichment screening using SELEX technology and next-generation sequencing (NGS), as shown in **Fig. 1a**. After data preprocessing, 1,082,299, 9,350, and 4,867 aptamer sequences of 35 nucleotides in length were obtained from each round. The model was initially trained with the sequence data from the third round and subsequently fine-tuned with the sequences from the sixth round to enhance its ability to identify enriched aptamer sequences, as depicted in **Fig. 1b**. We employed the byte-pair encoding (BPE) algorithm for sequence encoding during both training and fine-tuning phases. To evaluate the performance and generalization capabilities of the model, we divided the datasets into 90% for training and 10% for validation. Finally, we assessed the stability and binding ability of the Round 20 top-enriched aptamers and the screened candidates. By implementing these methodologies, we successfully developed a sophisticated language model named AptaGPT, which provided profound insights into aptamer biology and the capability to generate targeted sequences.

**Fig 1.**
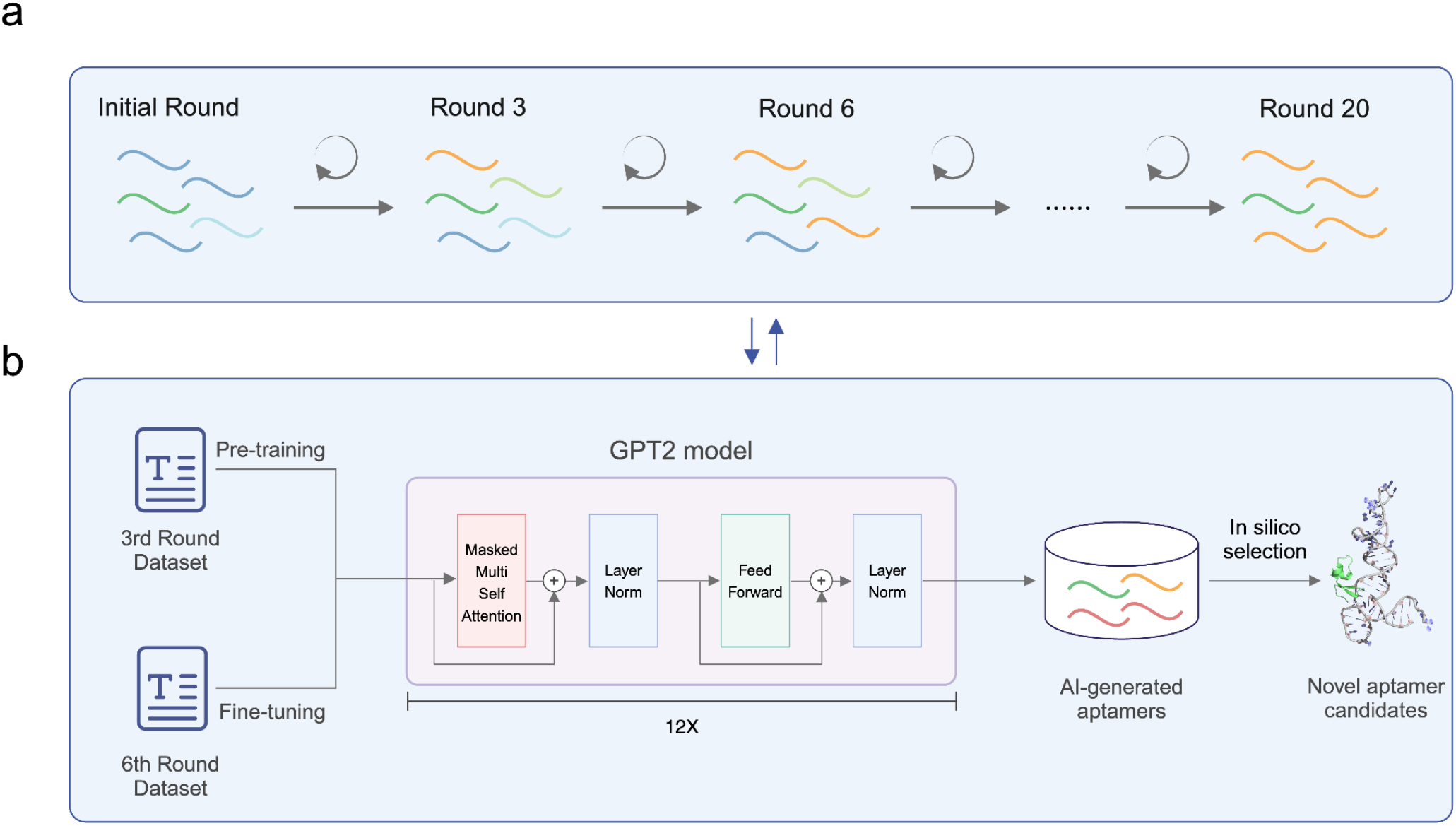
Overview of the workflow. **(a)** The SELEX process obtained aptamer sequences from the third, sixth, and twentieth rounds to construct the AptaGPT. **(b)** General workflow for designing aptamers using AptaGPT, incorporating in silico selection methods.

### AptaGPT enables aptamer design

In autoregressive language generation, the process of generating sequences involves iterative computation of conditional probability distributions to facilitate the step-by-step prediction and assembly of new words within a sequence. A primary focus of current research is identifying an optimal decoding approach for producing coherent and diverse textual output. Although the conventional greedy search method is computationally simple, it tends to yield repetitive text because it consistently selects the most common term at each time step. By contrast, beam search mitigates this problem by considering multiple potential words that are more likely to be correct. However, the text generated by beam search may still be repetitive and lacks originality^22^. Conversely, stochastic sampling methods, such as the top-kbased sampling strategy, randomly select a word from a set of k words with the highest probabilities^23,24^. This method better mimics the natural language generation process and is widely used in protein design^20,21^.

Therefore, AptaGPT generated aptamer sequences via a random top-k sampling method. The most suitable value for the k parameter was identified by evaluating the naturalness of the sequences based on the Jaccard index, a statistical measure utilized for assessing the similarities and differences among sets of samples. Examination of various k values ranging from 50 to 1000, as illustrated in **Supplementary Fig. 2a**, revealed that the highest score of 0.295 was achieved at k=700. Furthermore, the repetition penalty parameter was adjusted to 1.2, following recommendations from other generative models^20,25^, to improve the quality of the sampling outcomes. In the next step, AptaGPT performed ten iterations of sequence generation according to the specified parameters, returning one thousand sequences per iteration, for a total of ten thousand sequences, most of which were 35 nucleotides in length (**Supplementary Fig. 2b**). Some sequences were found to have a repetition frequency (**Supplementary Fig. 2c**). After selecting sequences of 35 nucleotides in length and excluding erroneous sequences, a total of 8,847 unique aptamer sequences were retained for follow-up analysis.

**Fig 2.**
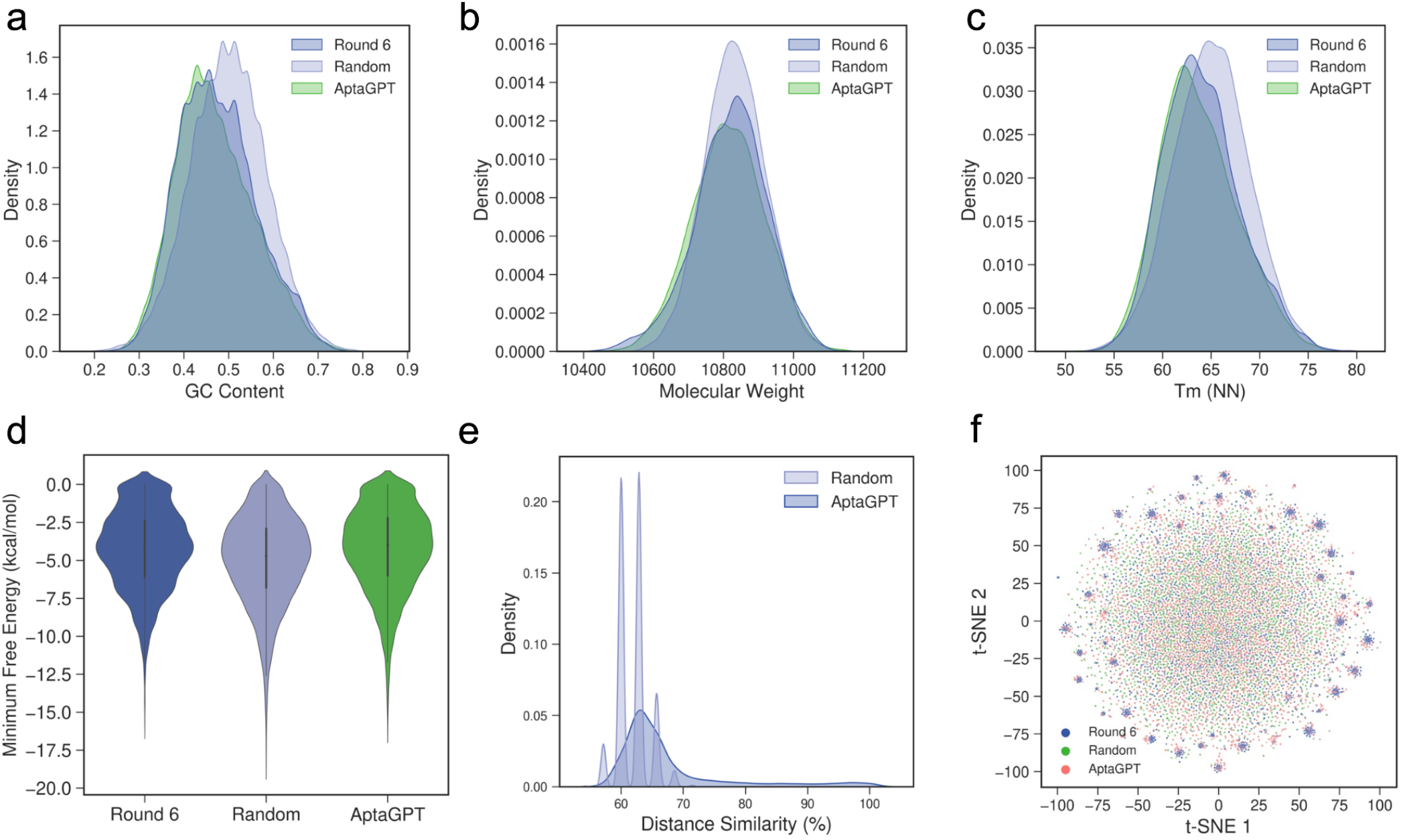
Evaluation of aptamer sequences generated by AptaGPT. **(a)** Distribution of GC content for Round 6, Random, and AptaGPT. **(b)** Distribution of molecular weight for Round 6, Random, and AptaGPT. **(c)** Distribution of nearest-neighbor melting temperature (Tm_NN) for Round 6, Random, and AptaGPT. **(d)** Distribution of the minimum free energy between the three groups of Round 6, Random and AptaGPT. **(e)** Distance similarity of AptaGPT and Random sequences to Round 6 sequences. **(f)** t-SNE distribution of Round 6, Random, and AptaGPT sequences after numerical encoding.

### AptaGPT generates diverse sequences

In order to assess the quality of AptaGPT-generated aptamers, several computational methods were used to compare the similarities between the sequences obtained from Round 6 and those generated by AptaGPT. In addition, we generated 10,000 35nucleotide sequences composed of four nucleotides to evaluate the quality of the AptaGPT-generated sequences against random sequences. We initially assessed the similarity among the three sets of sequences based on DNA physicochemical properties, such as GC content (**Fig. 2a**), molecular weight (**Fig. 2b**), and nearest-neighbor melting temperature (Tm_NN) (**Fig. 2c**). The results show that random sequences follow a normal distribution in these properties. In contrast, AptaGPT is more similar to Round 6. As illustrated in **Fig. 2d**, we employed the RNAfold program to determine the minimum free energy of the secondary structure. While random sequences exhibited lower minimum free energies, the energy distribution of sequences generated by AptaGPT closely approximated that of the Round 6 sequences. This suggests that AptaGPT can effectively replicate the structural features of actual aptamer sequences. Furthermore, we calculated the Levenshtein distances between random sequences, AptaGPT sequences, and Round 6 sequences to determine the maximum similarity. As depicted in **Fig. 2e**, the AptaGPT sequences exhibit over 80% similarity coverage compared to the random sequence. Moreover, the similarity between the AptaGPT sequences and the Round 6 sequences predominantly falls within the 60% to 70% range. The results of motif clustering using MEME indicate that the sequences generated by AptaGPT exhibit a similar pattern to that observed in Round 6 (**Supplementary Fig. 3**). Moreover, the visualization of the numerical encoding of the aptamer sequences demonstrates that the AptaGPT-derived sequences exhibit a comparable pattern to that of Round 6, significantly covering the outer sequence space (**Fig. 2f**). These findings indicate that AptaGPT enhances the diversity of generated sequences, thereby expanding the potential for identifying optimal candidates.

### In silico screening novel aptamers from generated space

In this study, we employed AptaGPT to generate a novel sequence space. It is essential to ascertain whether novel candidates can be identified from the generated sequences. To investigate this hypothesis, we developed a computational screening pipeline, as illustrated in **Fig. 3a**. In this selection process, we removed 133 identified sequences from Round 6 and created an initial library of 8,714 aptamers. The first stage of our filtering process utilized a G4RNA screener to determine the potential G-quadruplexes. G4-forming aptamers exhibit many advantages, including high stability, increased cellular uptake, ease of chemical modification, and low production cost^26,27^. The program’s prediction scores range from 0 to 1, with higher scores indicating a higher probability. We selected 246 aptamers using a threshold of 0.5. Subsequently, 153 aptamers were identified. Their minimum free energy, as predicted by RNAfold, was less than -5 kcal/mol. The distribution of these aptamers based on the G4 prediction score and minimum free energy is illustrated in **Fig. 3b**. Also, the motifs found in aptamers play a crucial role in their function by contributing to structural stability and specific binding. In this study, we utilized the FSBC tool to pinpoint and cluster the key motifs in the aptamer. As illustrated in **Fig. 3c**, we found the top three motifs, “AATGCA,” “TGCAG,” and “GCAGG,” to a filtering process, resulting in the identification of 82 aptamers. The last phase of our screening process entailed performing a molecular docking simulation to assess the strength and specificity of the interaction between the aptamers and the extracellular domain of BCMA. During this phase, we used the top 10 enriched aptamers from Round 20 as a reference, which had a median docking free energy of -235.08 kcal/mol. In contrast, the median docking free energy of the 82 candidate-generated aptamers was -241.53 kcal/mol (**Fig. 3d**). Among these candidates, six aptamers (A012, A024, A047, A048, A051, and A079) were selected for further analysis. Their interaction types with BCMA are shown in **Fig. 3e**, and the complex structures are provided in **Supplementary Fig. 4**.

**Fig 3.**
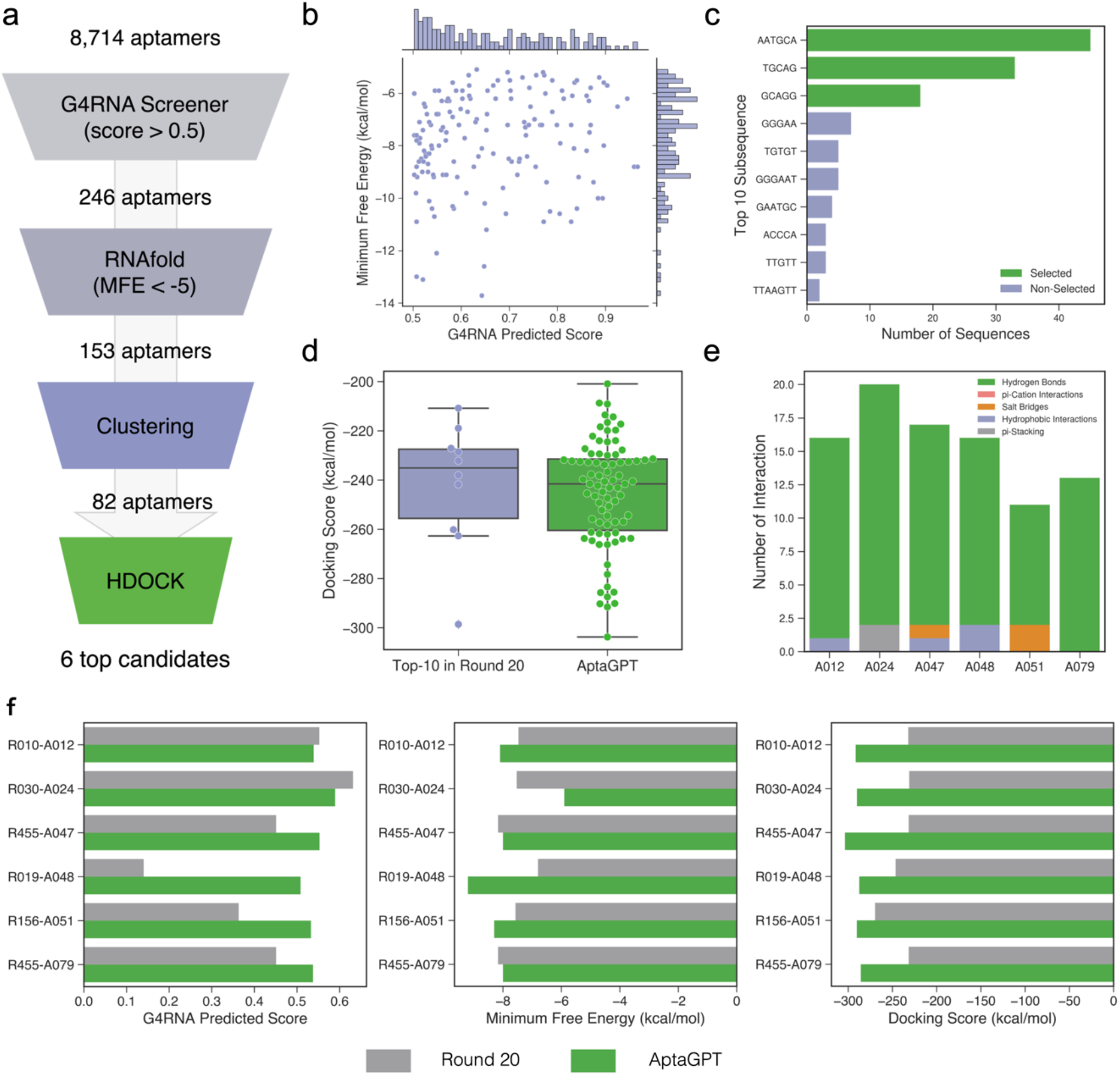
Computational screening of generated aptamer. (**a**) Flowchart of the in silico screening process, detailing the steps from the initial aptamer pool to the final selection of the top 6 candidates. (**b**) Distribution of G4RNA predicted scores and minimum free energies of secondary structures for 153 screened aptamers. (**c**) Rank of the subsequence identified by the FSBC algorithm for aptamer clustering. (**d**) Box plot of docking free energy of top-10 aptamers in Round 20 and selected aptamers. (**e**) Interaction types between the six candidates and BCMA. (**f**) Comparison of six candidate aptamers with their most similar sequences in top-1000 of Round 20 in terms of G4RNA prediction scores, minimum free energy of secondary structure, and docking scores.

To further evaluate the properties of these aptamers, the aptamers with the highest alignment identity to the six candidates from the top 1000 aptamers in Round 20 were screened, and their performance in structural and molecular docking was compared. The detailed results are listed in **Table 1**. As illustrated in **Fig. 3f**, the selected aptamers A047, A048, A051, and A079 exhibited enhanced G4RNA prediction scores, and A048 demonstrated improvement in minimum free energy of secondary structure compared to R019. Regarding docking free energy, A012, A047, and A051 exhibited more significant than 90% identity to the round 20 sequences, yet their free energies were significantly lower. The other three generated aptamer candidates also demonstrated promising results. These findings indicate that the aptamer mutants generated via AptaGPT may exhibit enhanced binding ability and structural stability compared to the subsequent SELEX-enriched sequences.

**Table 1.**
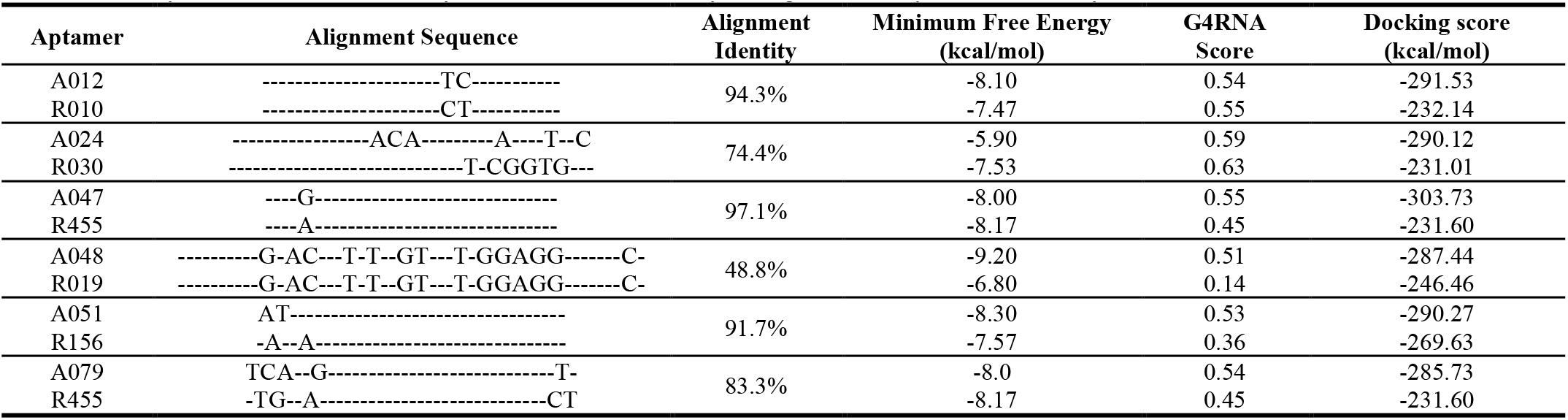
A comparison of six candidate aptamers with their corresponding similar sequences in the top 1000 of Round 20.

### Structure analysis of docking complexes

The AptaGPT model found two aptamers, A012 and A047, that showed significantly improved binding abilities compared to their Round 20 counterparts, namely R010 and R455, differing by one and two base units, respectively. These minor base changes greatly enhanced the interaction with BCMA by altering the spatial conformation of the aptamers. **Figure 4a** demonstrates that the two nucleotide variations of A012 resulted in remarkable alterations to its three-dimensional structure. This allowed A012 to engage in binding interactions with BCMA through 13 binding residues, which is three residues more than in R010. As a result, the binding stability and specificity of A012 may be improved. In A047, the nucleotide at the fifth position underwent a mutation, changing from adenine (A) to guanine (G) (**Table 1**). This slight modification has enhanced the interaction interface between the aptamer and BCMA. By optimizing the structure, we were able to increase the contact area between the two (**Fig. 4b**). This expansion of the binding pocket from nine to fifteen residues for R455 may significantly improve the affinity and selectivity of the interaction. In conclusion, these results show that even minor changes to the nucleotides play a crucial role in altering the structure and improving the function of aptamers. This also demonstrates the potential and usefulness of AptaGPT in aptamer design.

**Fig 4.**
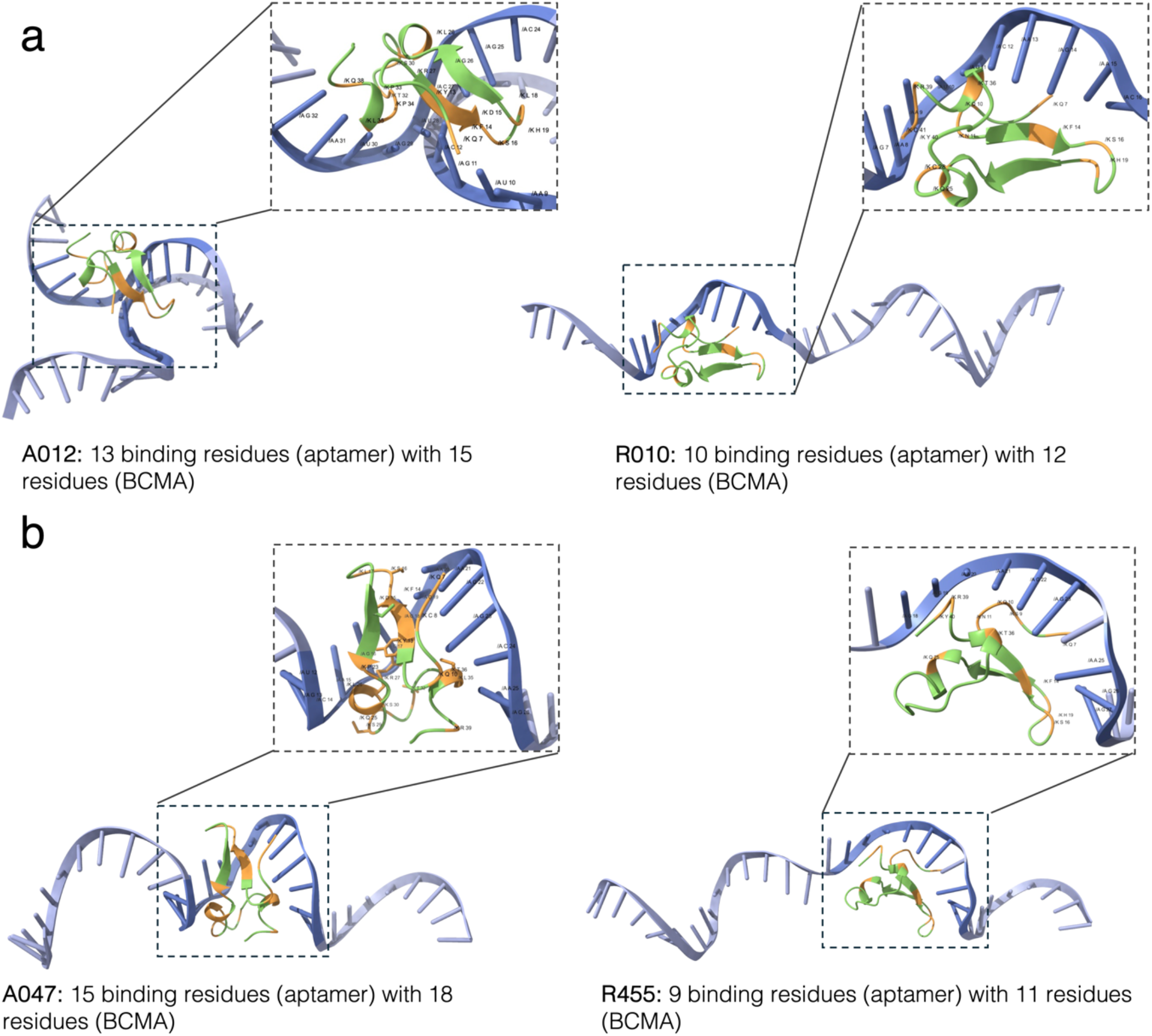
Comparative docking complexes of aptamer-BCMA interactions. **(a)** A comparison of the docking complexes between aptamers A012 (from AptaGPT) and R010 (from Round 20) with BCMA, focusing on the interacting residues. **(b)** A comparison of the docking complexes between aptamers A047 (from AptaGPT) and R455 (from Round 20) with BCMA, with a detailed analysis of the interaction sites.

## Discussion

The development and application of AptaGPT mark a significant advancement in aptamer research. By integrating a Generative Pre-trained Transformer (GPT) model with SELEX data, we have demonstrated the potential to streamline the aptamer discovery process, which traditionally has been a bottleneck in the field due to its time-consuming nature. The AptaGPT model’s capability to produce a wide range of high-affinity aptamer sequences demonstrates the potential of machine learning in enhancing computational biology.

One of the most striking findings from our study is the model’s ability to extend the diversity of sequences found in the later rounds of SELEX. This suggests that AptaGPT is not only learning the sequence patterns that are enriched through the SELEX process but also can explore new sequence spaces that have yet to be physically tested. The overlap in sequence motifs between the sequences generated by AptaGPT and those from round 6 of SELEX, as well as the differences in sequence similarity, highlight the ability of the model to capture evolutionary information in the sequences.

The in silico screening process further illustrates the practical utility of AptaGPT. By employing a multi-stage filtering approach, we could narrow the vast library of generated sequences to a select few that showed promising binding affinities in molecular docking simulations. Identifying six candidate aptamers with docking scores indicative of binding solid interactions with the target BCMA is particularly encouraging. These candidates serve as proof of concept for the efficacy of AptaGPT and as potential leads for further experimental validation.

However, it is essential to acknowledge the limitations of our study. While the AptaGPT model has shown promise in generating sequences similar to those enriched in SELEX, the actual test of its utility will be in the experimental validation of its predicted aptamers. Additionally, the molecular docking scores, while useful for screening, are not definitive predictors of binding affinity and specificity. Experimental assays will be necessary to confirm the binding characteristics of the selected aptamers.

Future work will focus on the experimental validation of the AptaGPT-selected aptamers and the refinement of the model’s predictive capabilities. Integrating additional data from SELEX experiments, including those with different targets and conditions, will likely improve the model’s generalizability. Moreover, the application of AptaGPT to other classes of targets beyond BCMA could further demonstrate its versatility and impact on the field.

In conclusion, AptaGPT represents a novel and powerful tool for aptamer design, potentially accelerating the discovery process and expanding the repertoire of aptamers available for therapeutic and diagnostic applications. As the field of computational biology continues to evolve, the integration of advanced machine learning models like GPT with traditional biotechnological approaches is poised to drive innovation and enable breakthroughs in biomedical research.

## Materials and Methods

### Dataset

In this study, we generated a set of SELEX datasets against BCMA targets for testing. Specifically, in rounds 3, 6, and 20 of SELEX, we acquired 28,107,584, 8,905,960, and 8,691,400 initial sequences by NGS technology, respectively. In the initial stages of data processing, aptamer sequences were retained if they were of a length of 35 nucleotides. In the third round of screening, the relative diversity frequency (RPM) threshold was set to 0.05, resulting in the exclusion of 108,229,900 aptamers. To further enrich the target sequences, the RPM threshold was increased to 0.08 in the sixth round of screening. Additionally, sequences that were duplicated in the third round were excluded. This resulted in the acquisition of 9,350 sequences for subsequent model fine-tuning. In the 20th round of screening, 4,867 sequences were identified that exhibited a significant degree of enrichment.

### Vocabulary encoding

For vocabulary construction, we used the SentencePiece^28^ to encode aptamer sequences and evaluated two tokenization strategies: Byte Pair Encoding (BPE)^29^ and Unigram language model^30^. The performance of these approaches was evaluated over a range of vocabulary sizes, from 2^10^ to 2^15^, as shown in **Supplementary Fig. 1**. The experimental results suggest that using the BPE approach leads to a decrease in the number of tokens required as the vocabulary size increases. In contrast, the average length of the tokens eventually increases. In contrast, when using the unigram approach, especially in scenarios with larger vocabularies, there is a more significant decrease in the number of tokens, and the average length of each token exceeds that of BPE. These observations suggest that the BPE method is more efficient with small to medium vocabularies, while the Unigram method may be more effective with larger vocabularies. Given the sample size and the aptamer sequences’ length, we chose to apply the BPE approach to the processing of aptamer sequences using a vocabulary size of 1024. The selection was made based on BPE’s proven effectiveness and reliability in situations with smaller vocabularies, making it highly suitable for accurate and efficient sequence coding and analysis.

### Model architecture

In this study, we employed the GPT-2 model^31^, a pre-trained generative language model based on the Transformer architecture^32^. This model was specifically designed to enhance language generation capabilities. The model configuration includes 12 attention heads, 768 embedding dimensions, and 86,631,936 parameters. The core of GPT-2 is a multi-layer decoder-style Transformer architecture^33^, where each layer comprises a multi-head self-attention mechanism and a positionwise feed-forward network. The multi-head self-attention mechanism operates according to the following formula:

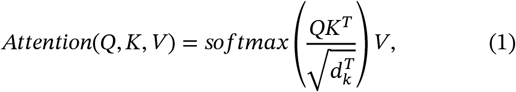

where *Q, K* and *V* represent the query, key, and value, respectively, and *d*_*k*_ is the dimension of the key vectors. This mechanism enables the model to be dynamically adjusted with each new word generated, thereby enhancing the relevance and coherence of the text based on the previous words.

The Hugging Face Transformers library was employed to facilitate the training and fine-tuning of the GPT-2 model (https://huggingface.co/openai-community/gpt2). In the pre-training phase, the model was trained for 20 epochs with a batch size of 256, a learning rate of 1e-4, a weight decay of 1e-2, and a maximum sequence length of 15. The model with the minimal evaluation loss was selected as the final model to ensure the effectiveness of the optimization results. In the fine-tuning phase, the model was adapted using a batch size of 128, a learning rate of 5e-5, and a weight decay of 0.01, with the number of epochs maintained at 20. The AdamW optimizer^34^ was used in both the training and fine-tuning phases.

### Sequence generation

In the task of generating aptamer sequences, we used the top k sampling method^23^ to control the randomness in the generation process and to improve the quality and relevance of the sequences. Top k sampling reduces uncertainty by considering only the top k most likely words from the model’s predicted vocabulary distribution. The math expression for this method is as follows:

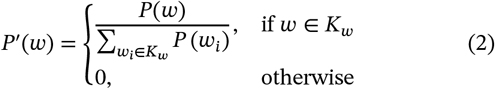

Here, *P*(*w*) represents the original probability distribution of the word *w*, and *K*_*w*_ consists of the top *K* words with the highest probabilities. To evaluate the quality and diversity of the generated sequences, we utilized the Jaccard index^35^. The Jaccard score is a metric for measuring the similarity between two sets, often used to assess the overlap of elements within sample collections. In our study, it is used to evaluate the overlap between Round 6 sequences and generated sequences. The calculation formula for the Jaccard score is:

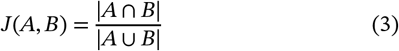

Where *A* and *B* are two sequence sets, |*A* ∩ *B*| represents the number of common elements between the two sets, and the total number of elements in the union of both sets. Our experiments tested different *k* values to determine the most appropriate one for our model and goals. K=700 was chosen as the optimal value, based on our goal of balancing diversity with sequence quality and similarity.

### Aptamer Sequence analysis

In our analysis, we employed the Biopython^36^ library to compute sequences’ GC content, molecular weight, and melting temperature and conducted pairwise alignment. To compare the similarity between random sequences, AptaGPT-generated sequences, and sequences from Round 6, we calculated the Levenshtein distance^37^ using the textdistance library. Motif clustering using MEME^38^ was used to identify conserved motifs or patterns in a set of aptamer sequences. During the vector encoding of aptamer sequences, numerical encodings were defined for the bases “A,” “T,” “C,” and “G” with values of 0.25, -0.25, 0.75, and -0.75, respectively. The t-SNE^39^ technique from the scikitlearn^40^ library was then applied for dimensionality reduction. In the clustering and screening process, we employed the FSBC algorithm^41^ for cluster analysis and identified vital motifs indicative of active regions in aptamers. Furthermore, we predicted the potential G-quadruplex scores in aptamer sequences using the G4RNA Screener ^42^. All analysis results were visualized using the matplotlib and seaborn libraries.

### Aptamer structure modeling

In this study, we used RNAfold and DRfold to predict the structure of the aptamer. The minimum free energy of the aptamer was determined using the RNAfold program at ViennaRNA^43^, and obtained its secondary structure. The tertiary structure was predicted using DRfold (https://github.com/leeyang/DRfold)^44^. DRfold is based on a combination of deep end-to-end analysis and geometric potentials. It allows high-resolution RNA structure modeling to be performed on a large scale.

### Molecular Docking

Molecular docking was performed using the HDOCKlite^45^ Linux program (http://huanglab.phys.hust.edu.cn/software/hdocklite). The extracellular region (residues 1-54) of human BCMA, derived from the Protein Data Bank (PDB) of 4ZFO^46^, served as the receptor. The predicted 3D structure of an aptamer was employed as the ligand. The receptor and ligand structures were inputted into the HDOCKlite to simulate and compute the docking free energy. The interaction types (e.g., hydrogen bonding, hydrophobic interactions, etc.) between the two were analyzed automatically by Protein-Ligand Interaction Profiler (PLIP, https://github.com/pharmai/plip)^47^. ChimeraX 1.7.1^48,49^ was used for the visualization of complex structures, and contacts were extracted using the default parameter, VDW overlap ≥ -0.4Å.

## Data availability

The SELEX sequencing data used in this study are not publicly available due to the ongoing research project’s needs. However, these data can be requested from the corresponding authors, subject to confidentiality agreements. The model weights are publicly accessible on the HuggingFace repository at the following link: https://huggingface.co/tmbj-aidd/aptagpt-bcma.

## Code availability

The model was trained using version 4.40.1 of the HuggingFace Transformers Trainer and version 2.3.0 of PyTorch. The training code and documentation can be accessed at the following links: https://huggingface.co/docs/transformers/main_classes/trainer.

## Supplementary Figures

**Supplementary Fig 1.**
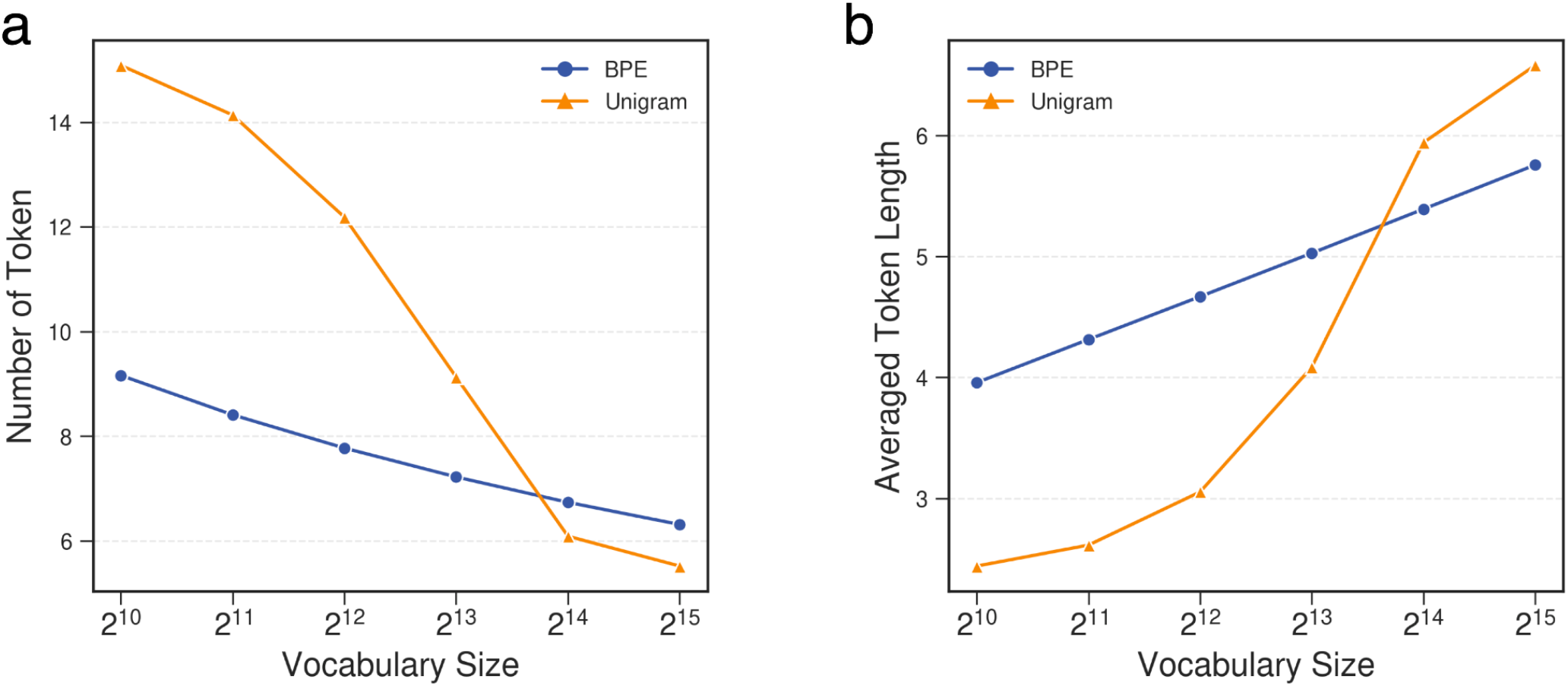
Comparison of Byte Pair Encoding (BPE) and Unigram tokenization at different vocabulary sizes. (**a**) Number of tokens generated by each method at different vocabulary sizes. (**b**) Average token length for each method with increasing vocabulary size.

**Supplementary Fig 2.**
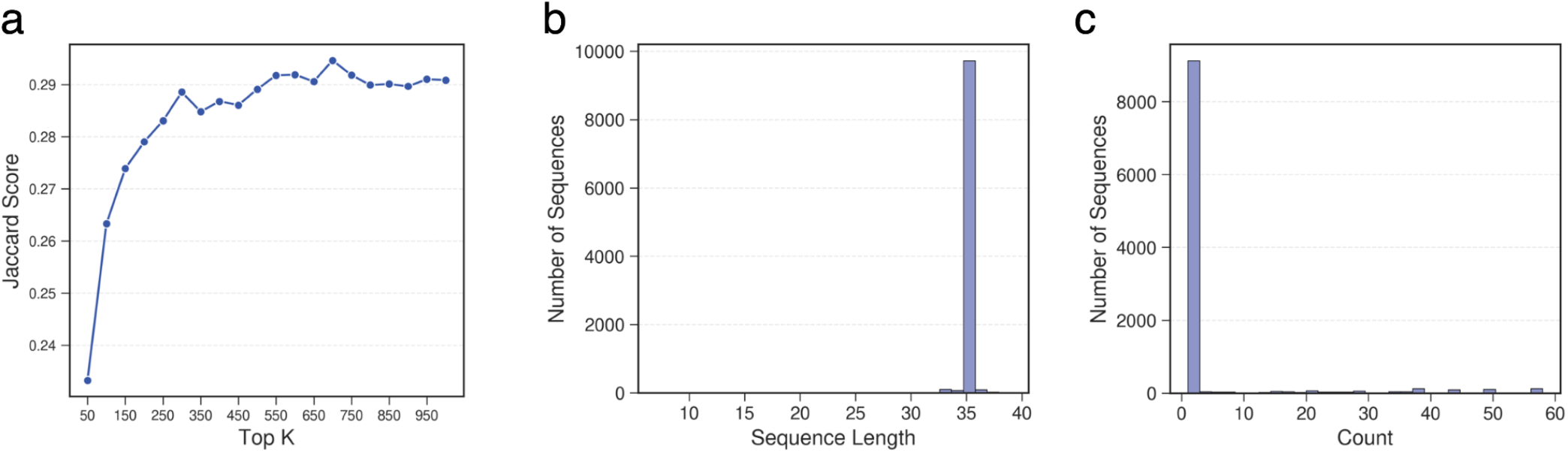
Sequence generation by AptaGPT. (**a**) Jaccard score of different top K values. (**b**) Sequence length distribution of the 10,000 generated aptamers. (**c**) Frequency distribution of unique aptamers.

**Supplementary Fig 3.**
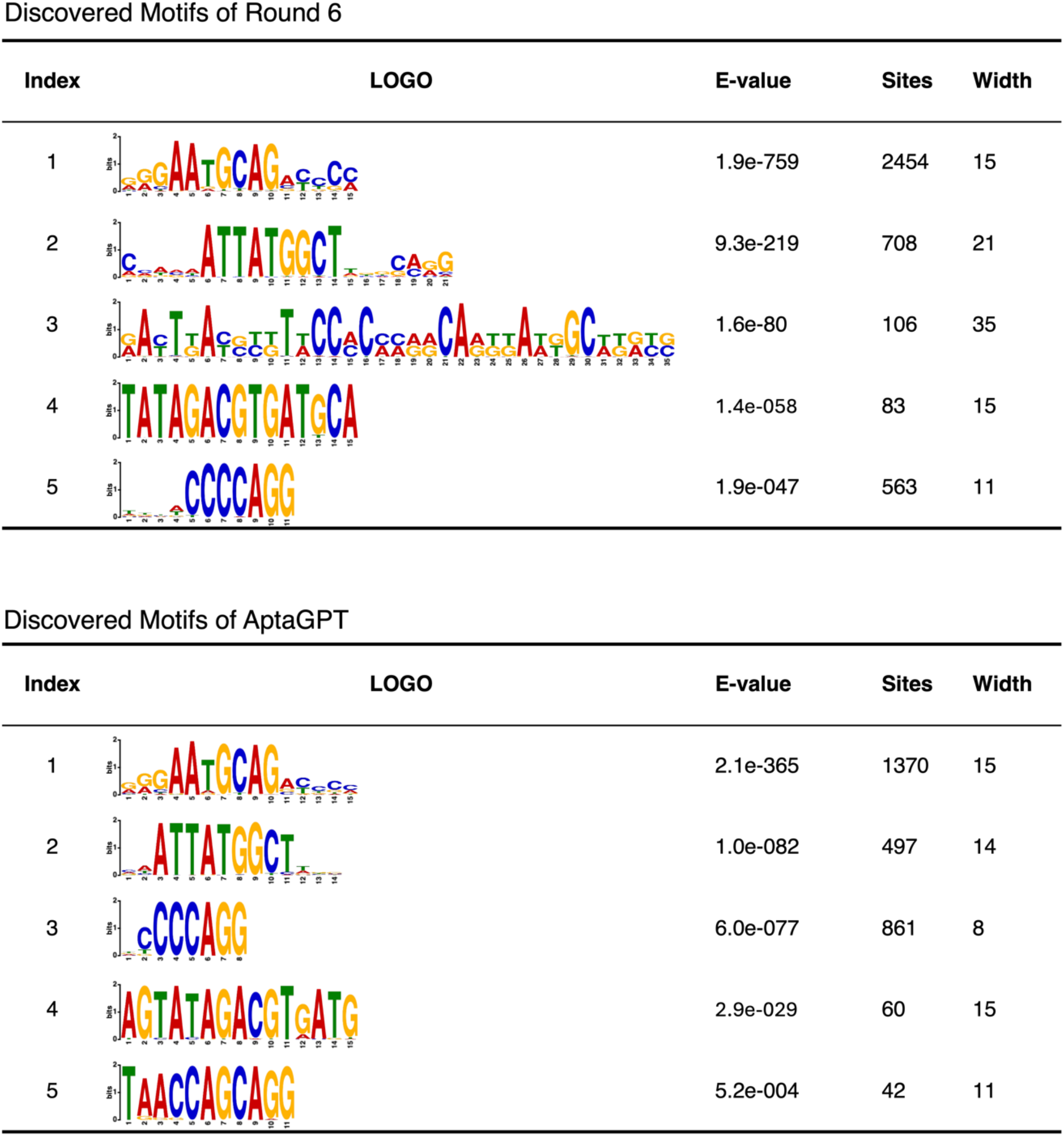
Clustering results of motifs analyzed by MEME. The sequence motifs identified in Round 6 and by the AptaGPT are illustrated as sequence logos. Each motif is ranked according to its E-value, which reflects its statistical significance. Additionally, the number of sites and motif width are included for each motif.

**Supplementary Fig 4.**
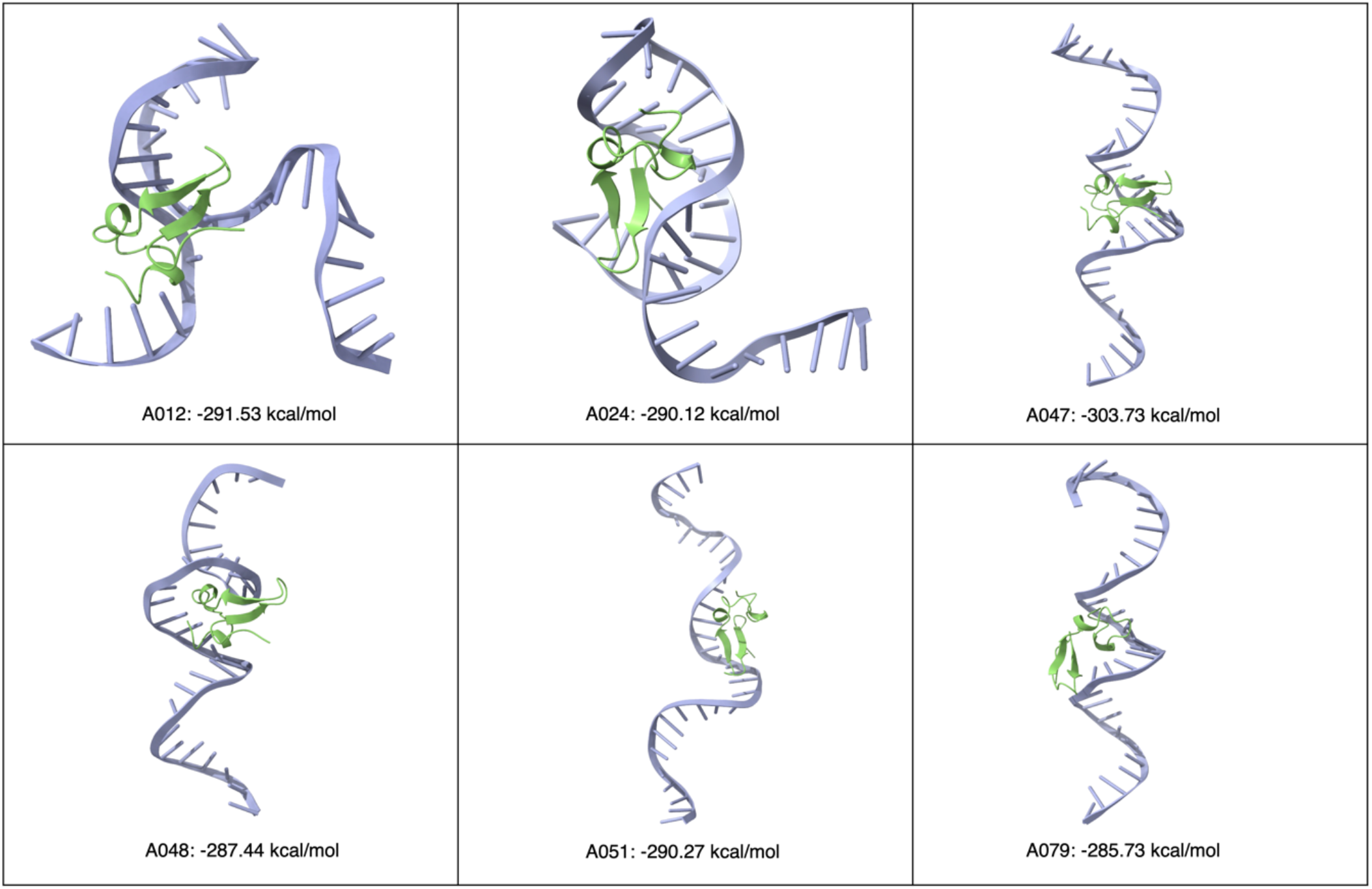
Predicted tertiary structures and docking complexes of the top six candidate aptamers. The predicted tertiary structures of the first six candidate aptamers using DRfold (shown in blue) and their docking complexes with the extracellular domain of BCMA obtained using Hdock are shown. The docking scores for each complex are in kcal/mol.

